# Self-assembly of polymer-encased lipid nanodiscs and membrane protein reconstitution

**DOI:** 10.1101/476556

**Authors:** Bikash R. Sahoo, Takuya Genjo, Kanhu C. Moharana, Ayyalusamy Ramamoorthy

## Abstract

The absence of detergent and curvature makes nanodiscs to be excellent membrane mimetics. The lack of structural and mechanistic model of polymer-encapsulated lipid-nanodiscs limits their use to study the structure, dynamics and function of membrane proteins. In this study, we parametrized and optimized the coarse-graining (CG) bead-mapping for two differently charged and functionalized copolymers, namely styrene-maleic acid (SMAEA) and polymethacrylate (PMAQA), for the *Martini* force-field framework and showed nanodisc formation (< 8 nm diameter) on a time scale of tens of microseconds using molecular dynamics (MD) simulation. Structural models of ~ 2.0 or 4.8 kDa PMAQA and ~2.2 kDa SMAEA polymer based lipid-nanodiscs highlights the importance of polymer chemical structure, size and polymer:lipid molar ratio in the optimization of nanodisc structure. The ideal spatial arrangement of polymers in nanodisc, nanodisc size and thermal stability obtained from our MD simulation correlates well with the experimental observations. The polymer-nanodisc were tested for the reconstitution of single-pass or multi-pass transmembrane proteins. We expect this study to be useful in the development of novel polymer based lipid-nanodiscs and for the structural studies of membrane proteins.

**TOC GRAPHICS:** 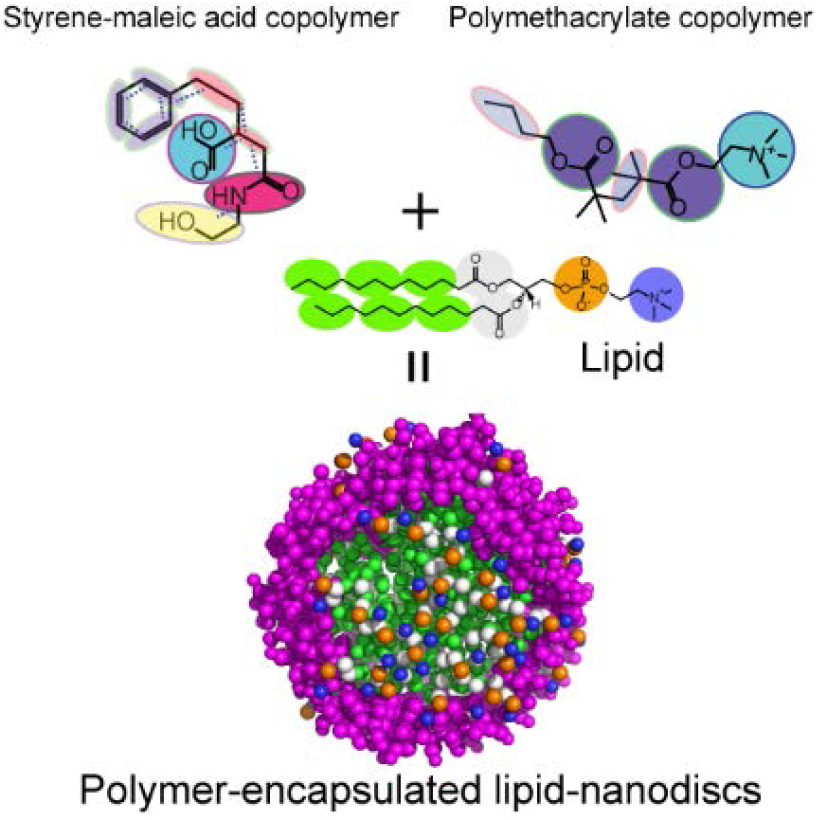

## 1. Introduction

Despite the recent advances in structural biology, membrane proteins continue to pose challenges to most biophysical techniques and biochemical approaches.^[1],[2]^ The aggregation kinetics of membrane proteins outside lipid membrane^[3]^ impulse researchers to develop methods to study their native structures^[4]^. As a result, reconstitution of membrane proteins using membrane mimetics such as bicelles, micelles, liposomes or nanodiscs^[4]^ has significantly eased the interrogation of membrane-associated proteins. Particularly, there is considerable interest in the development and application of nanodiscs^[5],[6]^. Use of nanodiscs as a potential chemical tool for protein misfolded diseases like Alzheimer’s disease is bound to create avenues for exciting biomedical applications^[7,8]^. These nanodiscs provide a detergent-free native-like lipid bilayer environment for functional reconstitution of a membrane protein or protein-protein complex. In addition, the size-tunable and lipid selectivity properties of nanodiscs have enabled researchers to design membrane protein selective nanodiscs for structural and functional studies^[9]^.

Recent studies have demonstrated the advantages of polymer based nanodiscs that include extraction of membrane proteins directly from cells, tolerance against pH and divalent metal ions, size tunability by simply changing lipid:polymer ratio, formation of macro-nanodiscs and their magnetic-alignment, and feasibility of applying biophysical techniques including solution and solid-state NMR experiments.^[10–13]^ The challenges and difficulties in membrane protein solubilization and purification has thus recently been overcome in native functional states by polymers. As an example, a maleic acid conjugated styrene/diisobutylene polymer recently showed stable and functional extraction of a human protein called rhomboid proteases.^[14]^ We have also demonstrated the advantages of the use of polymer based nanodiscs over peptide/protein based nanodiscs for the structural and functional investigation of amyloidogenic peptide interaction with lipid membrane at molecular level.^[15][16]^ In spite of the success, there are several disadvantages that need to be overcome for a wide spread application of polymer based nanodiscs. For example, the amphiphilic polymer has been shown to interact with the membrane protein of interest, which can interfere with folding and induce non-native conformational changes^[17]^. On the other hand, nanodisc forming polymers have recently been shown to have anti-amyloidogenic properties indicating their potential use in biomedical sciences.^[18]^ Synthetic modifications have been demonstrated to modulate the effect of polymer belt on the targeted protein structural and functional characterization. However, there is a lack of understanding of spontaneous nanodisc formation by copolymers at atomistic scale. Such information would enable the design of better suited polymer and polymer nanodisc for structural and functional investigation of a given membrane protein. While real-time monitoring of the self-assembly process to form nanodisc at atomic resolution is challenging, here we report a CG molecular dynamics (MD) simulation approach and demonstrate its use to understand the formation of nanodiscs. Specifically, parametrization and optimization of two functionally different copolymers, namely SMAEA and PMAQA, and their ability to self-assemble with lipids to form polymer nanodiscs are reported for the first time. Reconstitutions of three different membrane proteins including bacterial sensory rhodopsin (srII), amyloid precursor protein’s (APP) transmembrane domain and integrin-β3^[19–21]^ in these polymer nanodiscs are also demonstrated at atomic scale using microsecond timescale CG MD simulations.

## 2. Results and Discussion

### 2.1 Construction of polymer CG model

Parameterization and construction of CG models to perform MD simulation using SMAEA (2.2 kDa) and PMAQA (2.0 kDa or 4.8 kDa) polymers are shown in Figure 1 (a-b). CG bead mapping was accomplished based on experimentally identified functional groups (styrene, −N^+^R_3_, COO^−^) that are crucial for the nanodisc formation by the polymers.^[22],[23]^ The pseudo-bonds and bond lengths were determined from all-atom topology of the respective polymer (Figure. 1). Two differently sized PMAQA (~2.0 kDa and ~4.8 kDa) were tested with a similar CG bead mapping to understand the role of polymer length in the formation of nanodiscs. In addition, we employed two different lipids to evaluate the suitability of lipids for the nanodisc formation: zwitterionic lipid DLPC (1,2-dimyristoyl-sn-glycero-3-phosphocholine) and anionic lipid DLPS (1,2-dilauroyl-sn-glycero-3-phospho-L-serine).

**Figure 1.**
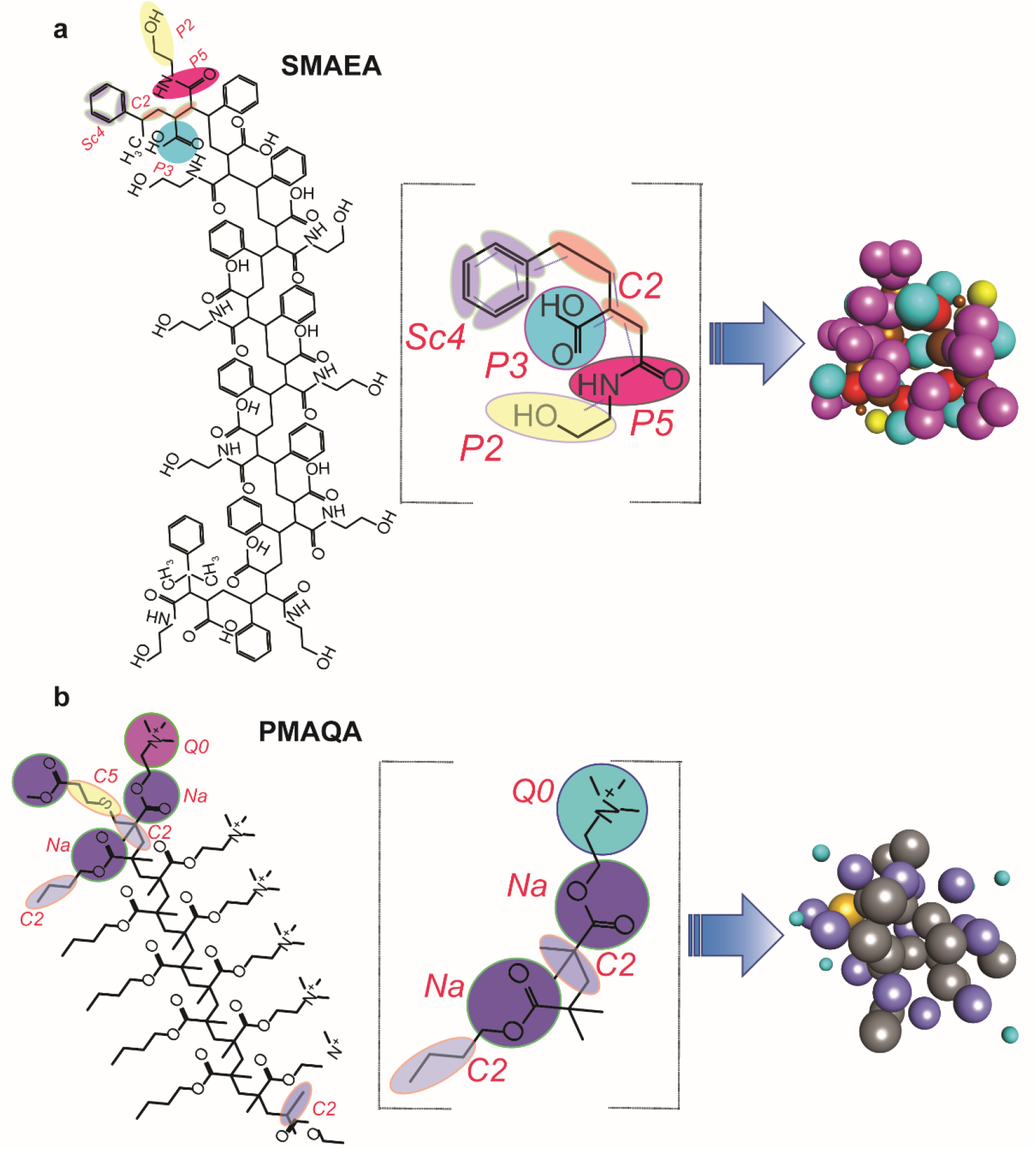
Coarse-grained (CG) mapping for the parameterization of SMAEA (a) and PMAQA (b) nanodisc forming copolymers. Chemical structures of SMAEA (top left) and PMAQA (bottom left) were built using *ChemDraw*. The selected CG groups are high-lighted in colors (middle column), and the CG bead mapping of each repeated units in SMAEA or PMAQA and their corresponding CG-models are shown in the right-most panel.

### 2.2 Solubilization of lipids by polymers

The solubilization of lipid-bilayers by a polymer has been shown to generate discoidal nanodiscs by light scattering, TIRF fluorescence and NMR experiments^[22,23]^. Under the experimental conditions, such morphological transition happens spontaneously and sometimes require mechanical procedures such as repeated freeze and thaw cycles depending on the lipid composition^[22],[23]^. Here, we investigate such a transition using a DLPC lipid bilayer model in Martini force field^[24]^. As illustrated in Figure S1, SMAEA was found to substantially destabilize the DLPC lipid bilayer (SMAEA:DLPC=1:4) over a time period of 4 μS MD simulation. The self-assembly of SMAEA on the DLPC lipid-bilayer facilitates membrane insertion and pore formation by SMAEA within ~ 2 μs. Segregation of upper and lower leaflet lipids with centrally bridged SMAEA were identified at the end of 4 μS MD simulation (Figure S1). Such membrane destabilization and transient pore formation by SMA copolymer (~7.4 kDa) was recently reported.^[25]^ Even though SMAEA disrupts DLPC lipid bilayer, we did not observe nanodisc formation within the microsecond timescale of MD simulation mentioned above. Therefore, these findings suggest that a longer timescale of MD simulation is essential to achieve sufficient disruption of the pre-structured lipid bilayer by the polymer to form a discoidal nanodisc as reported from experimental studies.

### 2.3 Self-assembly of polymers and lipids

In order to overcome the limitation mentioned above, we next performed MD simulation to monitor the spontaneous assembly of randomly distributed polymer (SMAEA or PMAQA) and free DLPC lipids (Figure S2) in solution at 1:4 polymer:DLPC molar ratio. Remarkably, the randomly distributed polymers and DLPC lipids in aqueous solution exhibited the formation of discoidal shape nanodiscs within 10 μs simulation (Figure 2a and b). The microsecond scale MD simulation showed formation of small-sized nanodiscs within several hundreds of nanoseconds followed by fusion to form a stable discoidal shape nanodisc in both polymers (Figure 2a and b). PMAQA and SMAEA copolymers with nearly equivalent size (~2.0 kDa), but differing in hydrophobic and hydrophilic functionalization, generated nanodiscs of size ~7.0 and 7.5 nm, respectively (Figure 2c and d). The polymers were found to be organized around the acyl chains of DLPC lipids exposing the lipid polar head groups to the solvent as reported from experimental observations.^[22],[23]^ The distribution of PMAQA, in comparison to SMAEA, surrounding the lipid tails were found to be distorted with few molecules localized on the membrane surface (Figure 2c). Atomistic inspection showed the N^+^R_3_ groups of the PMAQA polymer were located close to (~ 0.5 to 0.7 nm) the anionic phosphate groups of DLPC lipids (Figure 2c). A very strong hydrophobic lipid-polymer interaction is required to overcome the above-mentioned electrostatic polymer-lipid interaction (between N^+^R_3_ and PO4^−^ groups) to uniformly assemble all the polymer molecules around the acyl chains of the lipid bilayer; this can be accomplished by enhancing the hydrophobicity of PMAQA. In contrast, SMAEA distribution was found to be uniform like a belt around DLPC acyl-chains (Figure 2d, see Video SV1).

**Figure 2.**
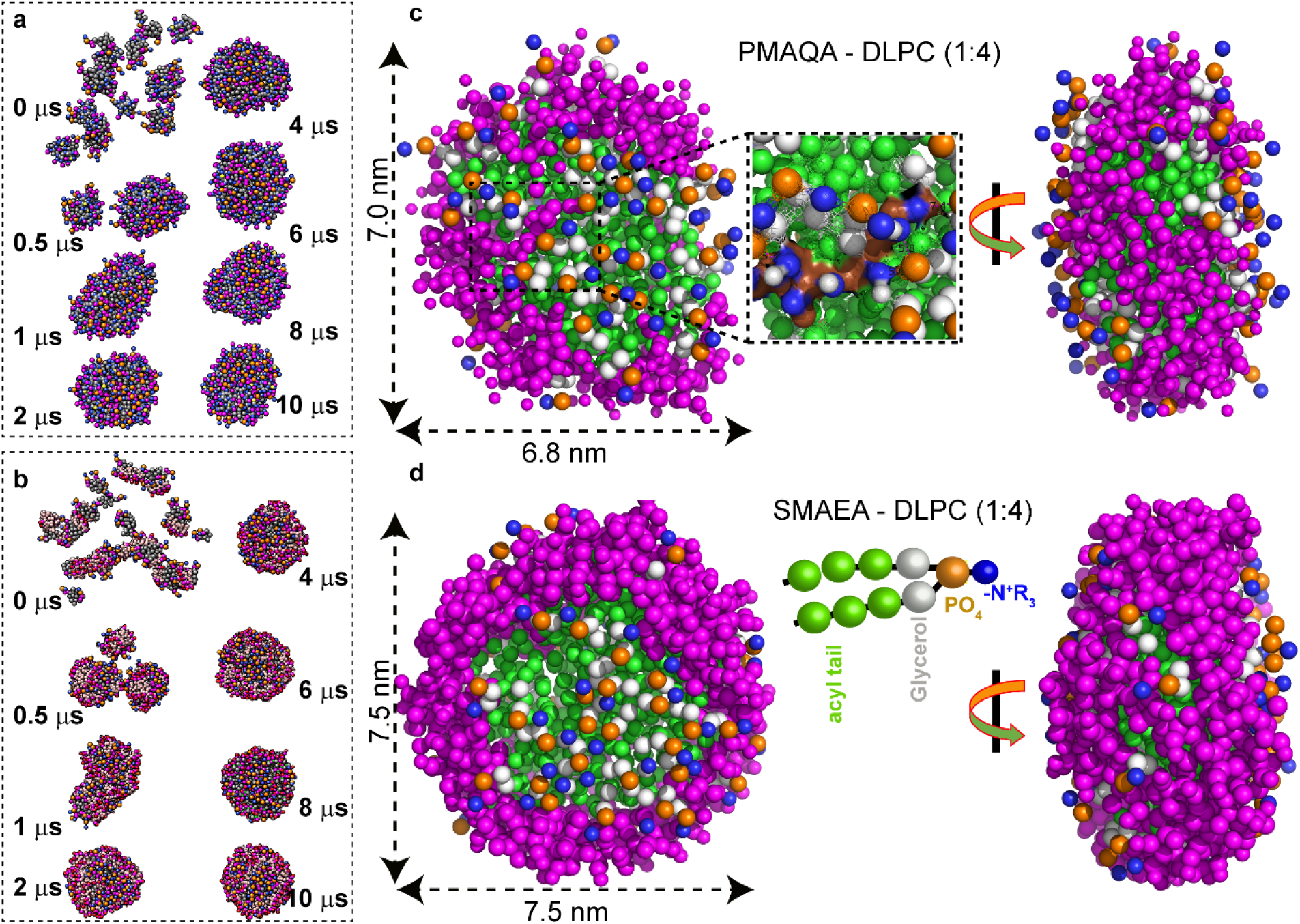
Coarse-grained MD simulation showing the spontaneous formation of DLPC nanodiscs on a time scale of 10 μs. Time-lapsed MD snapshots showing the self-assembling of (a) PMAQA-DLPC and (b) SMAEA-DLPC nanodiscs at 1:4 polymer:lipid molar ratio. Enlarged MD snapshots show the formation of discoidal DLPC nanodiscs of size ≈7 nm encased by PMAQA (c) or SMAEA (d); top-down (left) and side (right) views. The double-headed arrow indicates the average diameter of the nanodisc. The polymers are shown in violet and CG atoms of DLPC in other colors (c-d). Water molecules and ions are not shown for transparency.

### 2.4 Evaluation of lipid-bilayer properties in nanodiscs

Lipid bilayer thickness of a membrane mimetic plays a very important role on the membrane interaction, folding and topology of a membrane protein. Therefore, it is important to optimize the hydrophobic thickness of polymer based nanodiscs for a successful reconstitution of a membrane protein. While this is a daunting task to achieve experimentally, MD simulations can be used to test different parameters and provide guidelines for the design of a near-ideal polymer and optimized lipid composition to be used in a polymer nanodisc. Therefore, after successfully monitoring the self-assembly process underlying the formation of lipid nanodiscs for two different polymers as described above (Figure 2), MD simulation results were used to analyze the influence of the orientations of polymer molecules in a nanodisc on the encased lipid-bilayer’s thickness. As illustrated in Figure. 3a, SMAEA nanodisc exhibited a bilayer thickness of ≈3.5 nm at the center; whereas the boundary lipids surrounded by polymers depicted a low bilayer thickness. Such change in bilayer-thickness for nanodiscs has been observed in previous MD simulations for membrane scaffold protein (MSP) encased nanodiscs.^[26]^ Further estimation of bilayer thickness using a 20×20 matrix by GridMAT-MD presented an average bilayer thickness 2.57±0.13 nm which is in agreement with the experimentally determined hydrophobic bilayer thickness of 2.4 nm for DLPC (Figure. 3c).^[27,28]^ In addition, for a comparative analysis, we calculated the DLPC lipid bilayer thickness from a preassembled MSP-DLPC nanodisc generated using CHARMM_GUI^[29]^ and it was found to be 3.29±0.26 nm (Figure S3). On the other hand, the lipid bilayer surface distribution and random orientation of PMAQA molecules (Figure. 3b) exhibited a slightly lower average DLPC bilayer thickness of 2.08±0.24 nm (Figure. 3d). In a PMAQA-DLPC nanodisc, the thickness of several bilayer regions (blue regions in Figure 3d) was found to be <1.4 nm indicating a partial surface adsorption of PMAQA molecules (Figure 3b). A notable difference is the inhomogeneity of polymer-nanodiscs bilayer thickness when compared to the MSP-encased nanodiscs. This is due to the non-uniform stacking of polymer side-chains across the lipid acyl-chain unlike to the uniform and azimuthal alignment of the chains of two MSP proteins.^[26,30,31]^ But, similar to MSP nanodiscs, polymer nanodiscs showed an edge effect on bilayer thickness along with a thicker center bilayer region.^[30]^ However, an optimized membrane thickness for polymer nanodiscs may require longer time-scale of simulation to unify the edge effect of the polymer belt. The lateral diffusion rates of DLPC lipids calculated from the microsecond MD simulations were further compared between protein- and polymer-encased nanodiscs. The MSP-DLPC nanodisc presented a lateral diffusion rate of (4.37±0.78) × 10^−7^ cm^2^ s^−1^. The SMAEA- and PMAQA-DLPC nanodiscs showed a very little deviation in the lateral diffusion rates from that of MSP-DLPC with respective values of (4.33±0.61) × 10^−7^ and (4.43±0.15) × 10^−7^ cm^2^ s^−1^ at 303 K. The calculated diffusion rates for both polymer and MSP nanodiscs are of the same order magnitude (10^−7^ cm^2^ s^−1^) as observed experimentally^[32]^ and previous MD simulations.^[33]^

**Figure 3.**
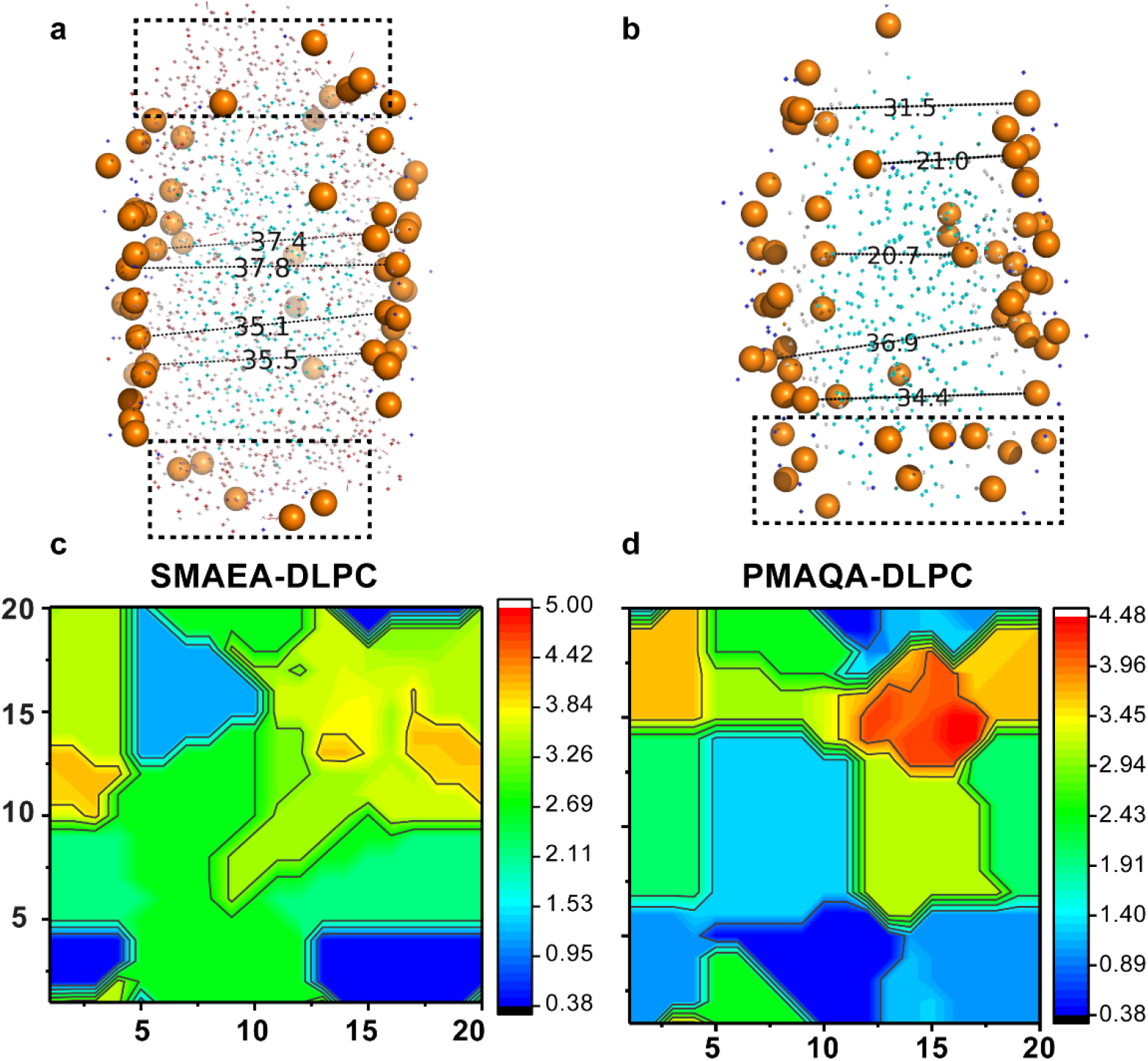
Lipid bilayer thickness in a polymer-encased DLPC nanodisc. PyMOL illustration of distance (in Å) between randomly selected upper and lower leaflet phosphate head groups in SMAEA (a) and PMAQA (b) nanodiscs retrieved at 10 μs MD simulation. The boxes in dashed lines represent the peripheral lipid heads close located to the proximity of polymers. The representative bilayer thickness is calculated using *GriDMAT-MD* from SMAEA-DLPC (c) and PMAQA-DLPC (d) nanodiscs and plotted in a 20×20 matrix. The scale bar shown is in nanometer.

### 2.5 Role of polymer concentration and lipid-specificity on nanodiscs formation

Since it is experimentally challenging to screen and optimize polymers and their lipid specificity to form stable nanodiscs, we examined the role of polymer’s molecular size using MD simulation. Experimental results showed that PMAQA with an optimal size of ~4.8 kDa forms nanodiscs.^[22]^ Here, we tested the nanodisc-forming capability of ~4.8 kDa PMAQA for a variable polymer:lipid molar ratio as shown in Figure 4. At 1:8 PMAQA:DLPC molar ratio, we observed spontaneous nanodisc formation with a uniform distribution of polymer molecules surrounding the hydrophobic acyl chains of DLPC lipids (Figure 4a). Atomic inspection revealed that the hydrophobic butyl groups (C2) of the polymer are oriented toward the hydrophobic lipid core region, whereas the cationic −N^+^R_3_ groups (Q0) of the polymer are exposed to the solvent (Figure 4a). The ~ 4.8 kDa PMAQA forms a DLPC nanodisc that is nearly equal to the size of SMAEA nanodisc (~ 7.5 nm diameter). The ~ 4.8 kDa PMAQA polymer nanodisc exhibited a lipid-bilayer with thickness of >3.5 nm for the central lipids (Figure 4c).

**Figure 4.**
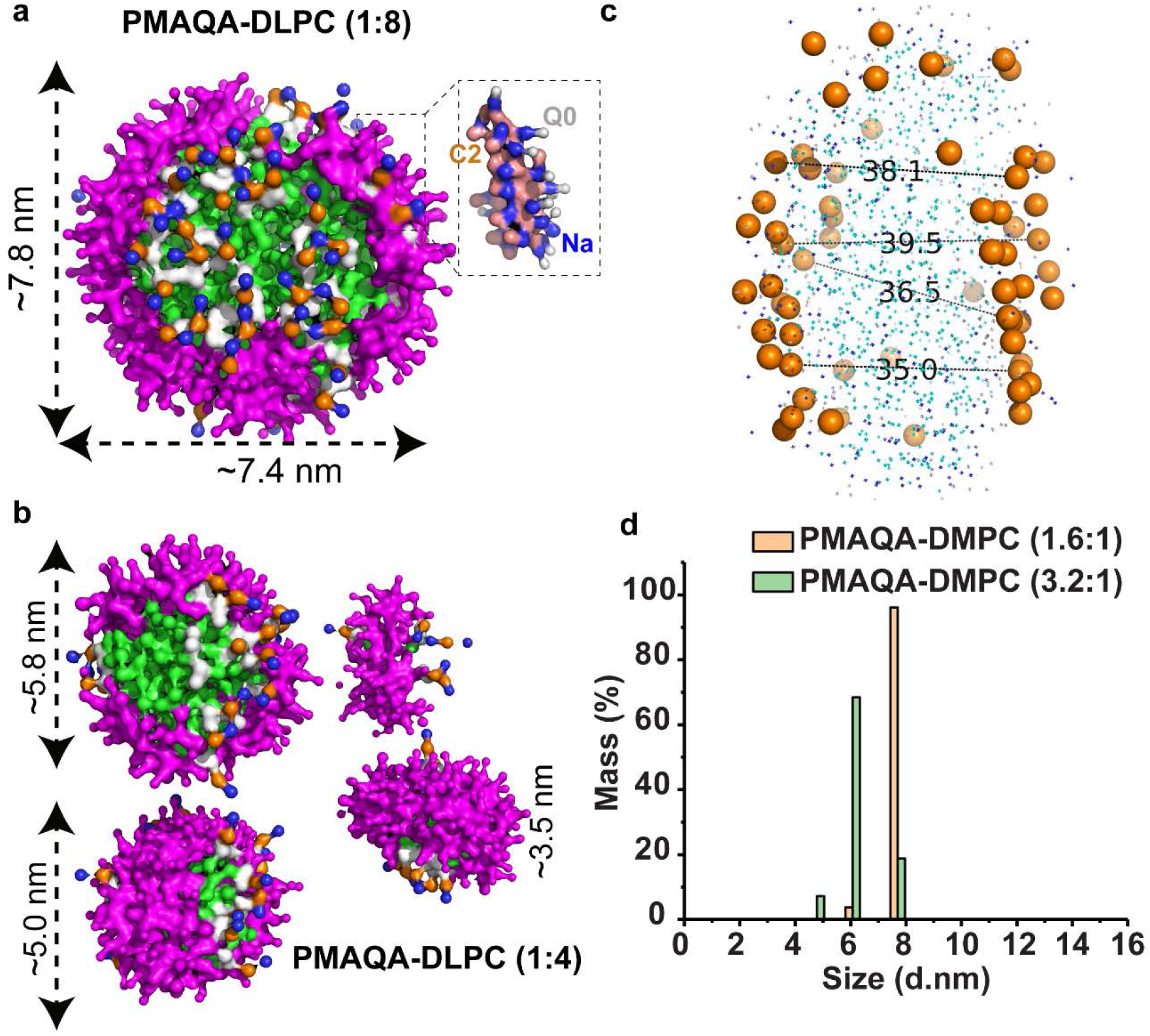
MD snapshots showing the formation of DLPC nanodisc by ~4.8 kDa PMAQA at a variable polymer to lipid molar ratio. Spontaneous formation of nanodiscs by ~ 4.8 kDa PMAQA at 1:8 (a) and 1:4 (b) polymer to DLPC molar ratio at 10 μs MD simulation. The polymers are shown in violet and DLPC atoms in different colors as indicated in Figure 2. Water molecules and ions are not shown for transparency. (c) The distance between the lipid head-groups (in Å) of PMAQA nanodisc shown in (a) are measured using PyMOL. (d) Dynamic light scattering showing the size distribution of PMAQA-DLPC nanodiscs at the indicated polymer to lipid molar ratio.

Next, we examined the effect of PMAQA polymer concentration on the size and morphology of nanodiscs formed. An increase in the concentration of PMAQA from 1:8 to a 1:4 molar ratio of PMAQA:DLPC resulted in smaller size nanodisc as expected from previously reported experimental results; in addition, nanodiscs with average diameters of ~ 5.8, 5.0 and 3.5 nm were also observed (Figure 4b). Unlike the PMAQA nanodiscs obtained from ~4.8 kDa at a 1:8 PMAQA:DLPC molar ratio (Figure 4a), the nanodiscs formed at higher concentration of PMAQA were found to contain PMAQA in random orientations and tightly packed (Figure 4b). Dynamic light scattering measurements of nanodiscs designed at a PMAQA:DMPC molar ratio corresponding to that used in MD simulations supported the computationally observed nanodisc size mentioned above (Figure 4d). As shown in Figure 4d, the ~4.8 kDa PMAQA at 1:8 polymer:DMPC molar ratio exhibited ~6.0 and ~ 7.7 nm size nanodiscs with a mass percentage of ~3.7 and 96.1%, respectively. On the other hand, for 1:4 PMAQA:DMPC molar ratio, ~ 4.7, 6.0 and ~ 7.7 nm size nanodiscs with a mass percentage of ~ 7.2, 68.4 and 18.8%, respectively, were obtained (Figure 4d).

MD simulation results (Figure 4) show that the lateral packing of lipids in PMAQA nanodiscs depends on the size and concentration of the polymer as expected from experimental results.^[22]^ The ability of MD simulations to reveal the location of different molecules constituting the nanodisc can be utilized in the optimization of experimental conditions to achieve ‘fluid lamellar phase’ like lipid packing which is crucial for functional reconstitution of a membrane protein or a protein-protein complex^[17]^. Since experimental studies have reported the difficulties in reconstituting various lipid composition in polymer nanodiscs,^[22],[23]^ it is important to examine the suitability of the lipid composition for a given polymer to form stable nanodiscs using MD simulations. For this purpose, we simulated the self-assembly of PMAQA or SMAEA polymers in presence of anionic DLPS lipids (with DLPC acyl chains). As shown in Figure S4, SMAEA forms a nanodisc within a time-scale of 10 μs, whereas PMAQA was found to be distributed on the membrane surface facilitated by its electrostatic interaction with negatively charged DLPS head groups.

**Figure 5.**
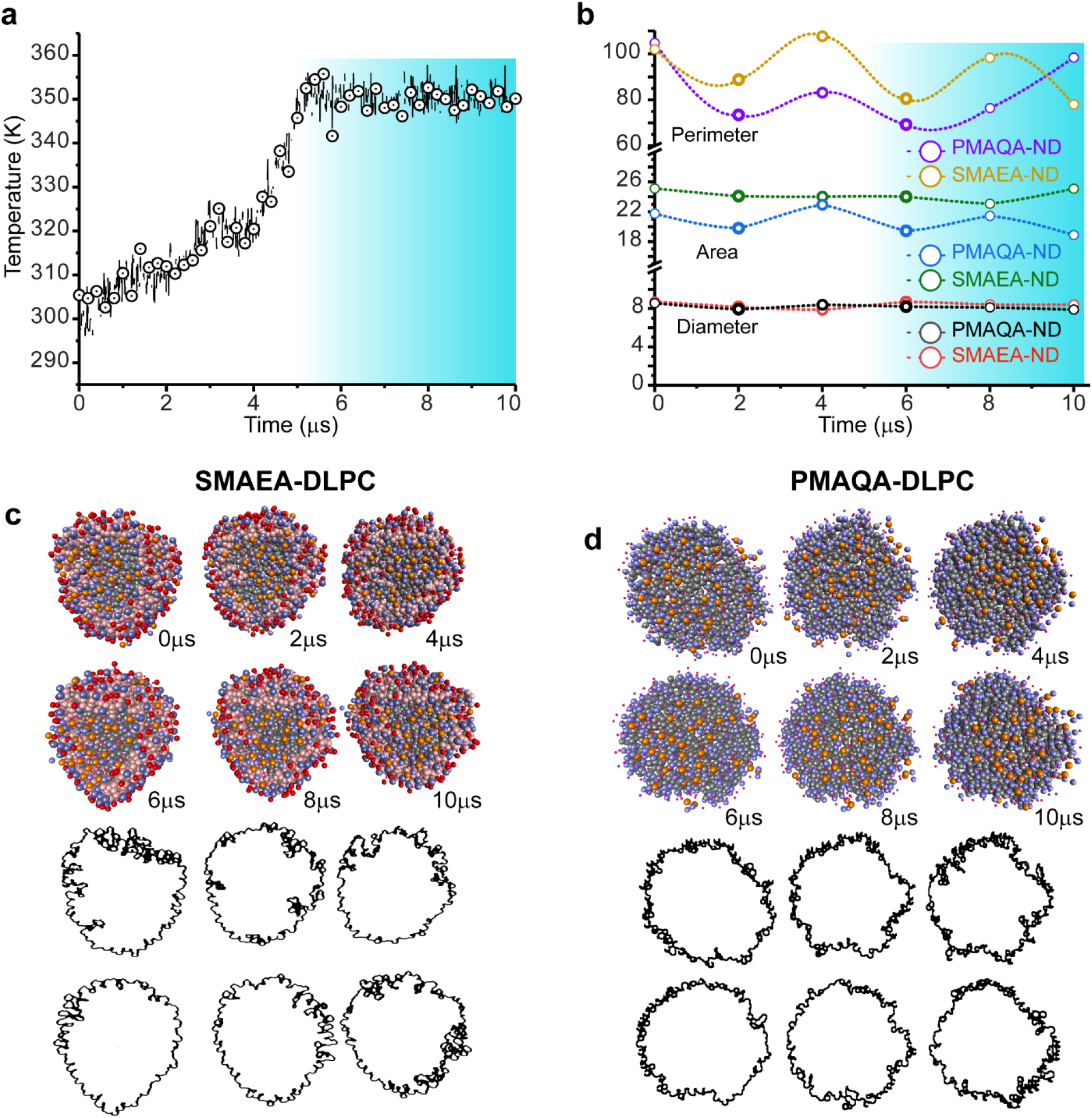
Thermal stability of SMAEA or PMAQA DLPC nanodiscs studied using simulated annealing MD simulation. (a) Polymer nanodiscs were subjected to a gradual increase in temperature from 298 to 415 K with respect to MD simulation time. (b) Feret’s diameter and area of nanodiscs calculated from MD snapshots retrieved from 1μs time interval using ImageJ. The highlighted regions in (a) and (b) show the variation in nanodisc size and shape with respect to temperature. (c and d) Illustration of polymer-nanodiscs analyzed using *ImageJ* (red) and the distribution of polymer-belt as a function of indicated simulation times (in μs). The scale for ImageJ analysis is shown in dashed line.

### 2.6 Thermal stability of polymer-nanodiscs

Thermal stability of nanodiscs is important for various applications including the biophysical and structural studies of membrane proteins^[34,35]^. Experimental studies have reported that peptide or MSP based nanodiscs can be stable up to ~ 60°C.^[36]^ Thermal unfolding has been seen with further increase in temperature that destabilizes the nanodisc’s size. In contrast to this, polymer-based nanodiscs are very stable and do not unfold even at 8? C.^[23]^ To better understand the thermal stability of polymer based nanodiscs, we investigated the thermal stability of PMAQA and SMAEA-DLPC based nanodiscs with respect to temperature increasing from 25 to ~80 °C as shown in Figure 5a. The simulated annealing experiment showed a minimal effect on the discoidal shape of polymer-nanodiscs with respect to temperature (Figure. 5c and d). Quantitative analysis of the polymer-nanodisc size showed an average Feret’s diameter of 8.17±0.26 nm for PMAQA (8.55 nm at t=0μs) and 8.37±0.32 nm for SMAEA-DLPC (7.87 nm at t=0μs) nanodiscs (Figure. 5b). Increase in temperature from 25 °C to 37 °C showed substantial rearrangement of the polymers around the lipid molecules as revealed from the change in area and perimeters of the nanodiscs (Figure. 5b). Further increase in temperature from 37 °C to 50 °C and 50 °C to 80 °C showed no significant change in Feret’s diameter; whereas periodic fluctuations in nanodiscs perimeter and area was noticed indicating the effect of temperature on both polymer and lipid spatial rearrangement. However, unlike the peptide or protein based nanodiscs, both PMAQA and SMAEA nanodiscs exhibited stability even at 80 °C as shown in Figure. 5c and d. An average area of 20.70±1.56 nm^2^ and 24.23±0.77 nm^2^ was calculated from MD snapshots taken at 2μs interval of time for PMAQA (21.74 nm^2^ at t=0 μs) and SMAEA-DLPC (24.02 nm^2^ at t=0 μs) nanodiscs, respectively (Figure. 5b). Overall, the simulated annealing MD simulation revealed the thermal stability of polymer nanodiscs up to 80 °C.

### 2.7 Reconstitution of transmembrane proteins in polymer-nanodiscs

Next, we studied the spontaneous reconstitution of a seven transmembrane domain bacterial sensory rhodopsin (srII) (PDB ID: 1XIO)^[37]^, and single transmembrane containing membrane proteins like integrin-β3 (PDB ID: 2L91)^[21]^ and amyloid precursor protein (APP) (PDB ID: 2LLM)^[38]^ in polymer nanodiscs. Membrane proteins were initially placed in different orientation such as ≈1 nm away from the SMEA-DLPC nanodisc or partially inserted as shown in Figure. 6 (top row). The srII protein was found to interact with the edge of the nanodisc (SMAEA belt) and the complex remained stable for several microseconds (~1 μs) followed by a change in the shape of the nanodisc from a discoidal to ellipsoidal shape (Figure S5). The lipid bilayer insertion of srII at 4 μs displaced several of SMAEA polymer molecules, and from 6 to 8 μs the seven transmembrane domains of the protein gradually oriented perpendicular to the plane of the lipid-bilayer and remain inserted until the end of the simulation (Figure S5). CG to all-atom conversion showed that the srII transmembrane domains were well oriented within the lipid-bilayer;^[19]^ however, the ~7.5 nm size nanodisc was found to be not efficient in maintaining the discoidal shaped structural integrity of SMAEA-DLPC nanodisc (Figure. 6a and S5).

We then studied the reconstitution of two different single-pass transmembrane proteins, namely APP and integrin-β3 (Figure 6b and c). Remarkably, unlike srII, APP and integrin-β3 were found to have little effect on the shape and size of SMAEA nanodiscs. Both APP and integrin-β3 were found to interact with nanodiscs within several hundreds of nanoseconds. The APP fragment that was marginally inserted into the nanodisc’s lipid-bilayer surface was found to incorporate its helix into the membrane and oriented at an angle ≈20° with respect to bilayer normal as observed previously^[39]^ (Figure 6b and S6). The N-(GSNK) and C-(KKK) terminal lysine residues were exposed to the solvent; whereas the centrally located helix residues were packed inside the nanodisc facing one side to the lipids and other to the polymer belt (Figure 6b). Similarly, integrin-β3 was found to localize in the lipid-bilayer with ≈30° tilt of the transmembrane helix with respect to the plane of lipid-bilayer, which is in agreement with previous reports that suggested an increase in the tilt angle in the absence of its other subunit’s TM domain (Figure 6c and S7).^[40,41]^

**Figure 6.**
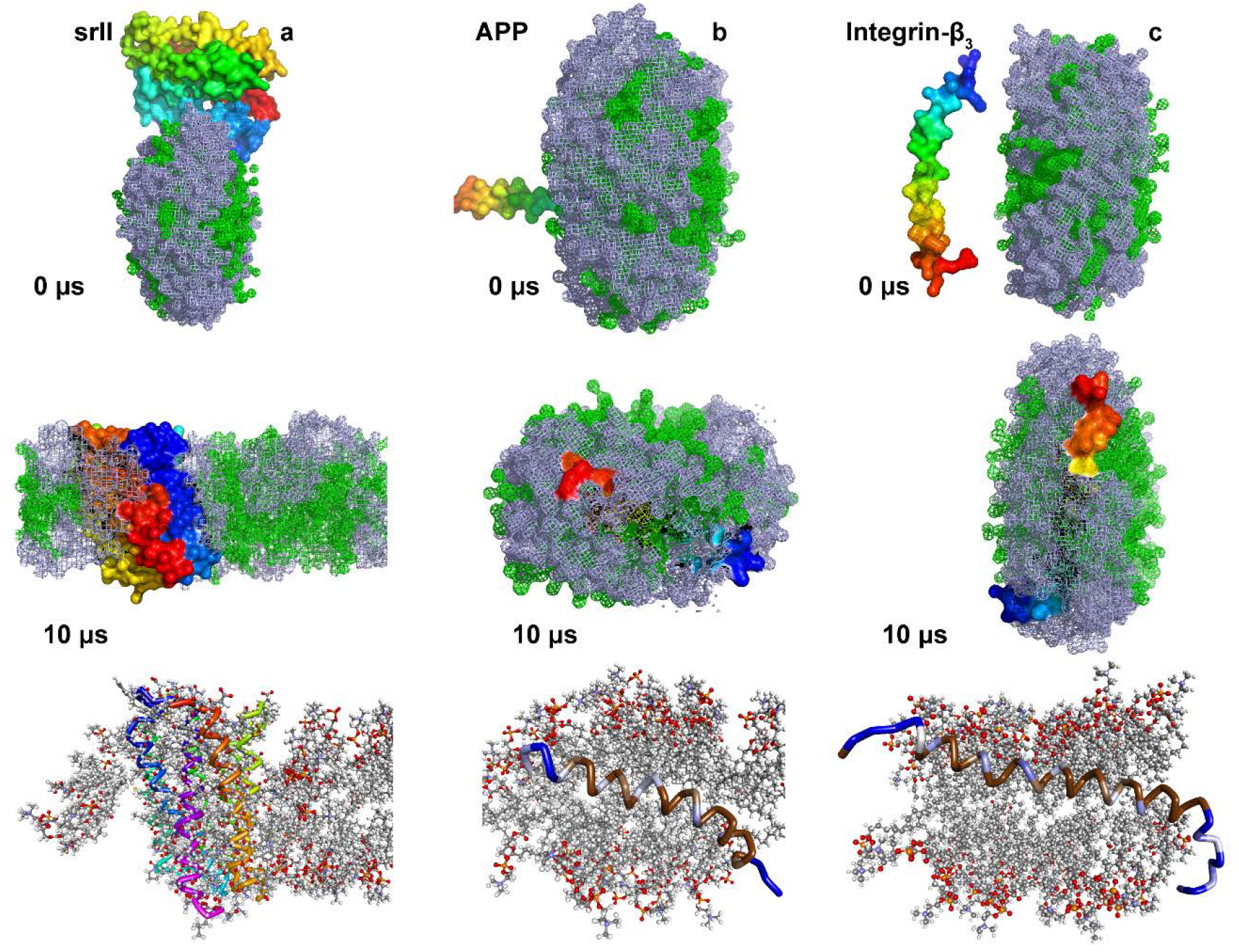
MD snapshots showing the reconstitution of membrane proteins in a SMAEA-DLPC nanodisc. Monitoring the reconstitution of seven transmembrane domain sensory rhodopsin-II (a), single-pass transmembrane amyloid-precursor protein (b) and integrin-β3 (c) into a SMAEA nanodisc (shown in Figure 2b) using 10 μs CG MD. Polymers (grey) and DLPC (green) are shown in mesh and proteins as surface in PyMOL with N-and C-termini in blue and red, respectively. The corresponding all-atom models generated from Martini CG model are shown in the bottom-most row. Membrane proteins are shown in ribbon and lipids in ball-stick. Zoomed structure of all-atom model structures are given in Supplementary Figures S6 and S7.

## 3. Conclusions

In conclusion, we have successfully demonstrated the formation of lipid-nanodiscs by two different amphiphilic copolymers consisting of styrene or polymethacrylate moieties using atomistic CG MD simulation. We expect that the parametrization of the polymers and methodology presented in this study will be useful in designing and screening new polymers that can efficiently form nanodiscs. The optimization of polymer length, lipid specificity and stability would not only provide a better understanding of the polymer-belt interference with the target membrane protein and its function, but also be useful in the development of nanomedicine or peptide membrane interaction studies as demonstrated in this work.^[15]^ We foresee that the computational approach employed here can be generalized by varying the polymer hydrophobic and hydrophilic moieties to design membrane protein selective nanodisc systems for a successful reconstitution. The coarse-grained parameters optimized for Martini force-field framework in this study for polymers will create avenues for further development of coarse-grained systems for other polymers to be used in biological and chemical studies.

## 4. Experimental Section

### 4.1 Chemicals

The PMAQA of ~4.7 kDa and SMAEA of ~2.2 kDa was synthesized and purified as reported elsewhere.^[15,22,23]^ 1,2-dimyristoyl-sn-glycero-3-phosphatidylcholine (DMPC) lipid was purchased from Avanti Polar Lipids, Inc^®^ (Alabaster, AL). All other reagents were purchased from Sigma-Aldrich^®^.

### 4.2 Dynamic light scattering

Large unilamellar vesicles (LUVs) of DMPC were prepared as described elsewhere.^[16,42]^ Briefly, DMPC lipids were dissolved in 1:1 chloroform and methanol and evaporated under the continuous steam of nitrogen gas and the lipid film was incubated o/n under vacuum. The DMPC lipid film was next hydrated in 10 mM sodium phosphate buffer (pH 7.4) followed by 5 minutes vortex mixing. The lipid mixture was suspended with a DMPC to PMAQA molar ratio of 1:1.6 or 1:3.2 followed by 5 minutes vortex mixing and incubated for 15 minutes at 37 °C under shaking. The mixed solution was subjected to several freeze-thaw cycles to homogenize the samples and were incubated overnight at 37 °C with gentle shaking to generate nanodiscs. Next, dynamic light scattering (DLS) measurement (Wyatt Technology Corporation) was performed to check the size distribution of PMAQA-DMPC nanodiscs using a 1 μl quartz cuvette at 25 °C.

### 4.3 PMAQA and SMAEA coarse-grained bead mapping

The 2D structure of PMAQA (~2.0 or 4.8 kDa) and SMAEA (~2.2 kDa) was generated using ChemBio Office Ultra 12.2 and exported to Chem3D 16.0. The chemical structures were energy minimized using MMFF94 force field^[43]^ for 10000 number of iterations with a minimum root mean square gradient of 0.1. The energy minimized structures were further subjected to molecular dynamics simulation in MMFF94 for 1,00,000 steps at 300 K. The all-atom topology of simulated structures of polymers were obtained by inputting the structures to the Automated Topology Builder^[44]^ and CCPN NMR AcpyPE.^[45]^ The all-atom 3D structure and topology files were considered for the parameterization of coarse-grained bead mapping in the Martini force field framework.^[46]^

The SMAEA polymer comprised of 9 repeating units of styrene, ethanolamine and carboxylic acid terminated with cumene. Each styrene group is represented with a three bead mapping (referred to tyrosine), ethanolamine with two bead mapping and carboxyl group with one bead mapping for the Martini framework as shown in Figure. 1. The bead selections were refereed from the standard Martini models (Martini 2.2P^[24]^) that has been tested for other biomacromolecules including proteins. The ~2 kDa PMAQA polymer comprised of 6 repeating units of quaternary ammonium (−N^+^R_3_) was assigned with one-bead referring to Martini phosphatidylcholine lipid topology. One-bead mapping for the hydrophobic butyl (C4) chain and ester groups was considered for the PMAQA CG model building. A similar CG bead mapping was considered for ~4.8 kDa PMAQA to test the role of polymer length in the formation of nanodisc. An optimal hydrophobicity (f) to hydrophilicity (1-f) fraction (f=0.5) was considered for PMAQA in reference to experimental results that showed nanodisc formation with “f” value ranging from ~0.3 to 0.6.^[22]^ The carboxyl group in SMAEA and quaternary ammonium group in PMAQA were assigned with one negative and positive charge, respectively.

### 4.4 MD simulation

The CG model for both PMAQA and SMAEA were generated from their all-atom MD simulation following the parametrizing of new molecule based on atomistic simulations documented in Martini (http://www.cgmartini.nl/index.php). Briefly, the polymers were simulated in aqueous solution with counter ions (Na^+^ or Cl^−^) for a production run of 100 ns using and the all-atom topology and trajectory files were used to generate CG MD simulation inputs in GROMACS version 5.0.7.^[47]^ A in-house python script was used to generate the indexing file of CG beads, angles and bonds by inputting the all-atom topology information and our CG bead mapping approach shown in Figure. 1.

To monitor the binding of polymer to lipid-bilayer, a CG MD system was built by placing SMAEA at a minimum distance ~1 nm from the DLPC lipid-bilayer in a cubic box system. The martini DLPC lipid was chosen for our study as it provide a general phosphatidylcholine lipid and correspond to atomistic C12:0 dilauroyl (DLPC)-C14:0 dimyristoyl (DMPC) tails. In our previous experimental study,^[22],[23]^ we demonstrated the spontaneous lipid-nanodisc formation by both polymers selectively for phosphatidylcholine lipid with C14 tails. The DLPC lipid-bilayer was built using Martini insane python program and simulated for 2 μs prior to SMAEA. The MD system was solvated using one bead water and neutralized by adding counter Na^+^ ions under a periodic boundary condition. The SMAEA-DLPC lipid bilayer (SMAEA to DLPC) MD system was next subjected to energy-minimization using the steepest-descent method followed by constant volume and pressure equilibration as described elsewhere^[48]^. The SMAEA-DLPC bilayer system was finally simulated for a production run of 10 μs at 303.15 K.

The CG models of spontaneous assembling of lipid and polymer was created by randomly placing the lipids and polymer at different molar ratio using *gmx insert.* The molar concentration of PMAQA or SMAEA were directly referred from our experimental findings to build the MD systems. A polymer to lipid (DLPC or DLPS) molar ratio of 1:4 was used for spontaneous MD simulation for ~2.0 kDa PMAQA and ~2.2 kDa SMAEA. 1:4 or 1:8 polymer to DLPC molar ratio was used for ~4.8 kDa PMAQA MD simulation. All MD systems were solvated using one-bead water model and neutralized using counter ions followed by energy minimization, equilibration and production run for a time-scale of 10 μs at 303.15 K. For comparative analysis of lipid properties, a CG model structure of MSP-DLPC nanodisc was built using CHARMM-GUI^[29]^ and simulated for 1 μs at 303.15 K.

The protein reconstitution MD systems were generated by placing the protein ~ 1 nm away from the SMAEA-DLPC nanodiscs obtained as the end product of spontaneous assembly MD simulation (at 10 μs). Three different membrane proteins such as bacterial rhodopsin (srII; PDB ID:1XIO)^[37]^, amyloid-precursor protein (PDB ID:2LLM; GSQKLVFFAEDVGSNKGAIIGLMV GGVVIATVIVITLVMLKKK)^[38]^ and integrin-β3 (PDB ID:2L91; PESPKGPDILVVLLSVMGA ILLIGLAPLLIWALLITIHDRKEF)^[21]^ with single or multiple transmembrane domains were considered for reconstitution MD simulation. The CG beads of targeted protein was generated using *martinize* python script. The protein-nanodisc MD systems were solvated using one-bead water model supplemented with 150 mM NaCl and the systems were neutralized with appropriate counter ions. Energy minimization, equilibration and production MD run of 10 μs at 303.15 K was performed to monitor the reconstitution of all three membrane proteins into SMAEA-DLPC nanodisc.

### 4.5 Bilayer thickness calculation

The lipid-bilayer thickness from polymer-encased lipid nanodiscs were calculated using GridMAT-MD program^[28]^. The end structure (at 10 μs) of each MD system was retrieved and subjected to GridMAT-MD to calculate the bilayer thickness in a 20×20 matrix and plotted using OriginPro (academic license). For comparative bilayer thickness analysis, the bilayer thickness of MSP-DLPC nanodisc built using CHARMM-GUI^[29]^ was calculated using GridMAT-MD. The protein and polymer-DLPC nanodisc lipid bilayer thickness was compared with each other and with the experimental values.^[27]^ The lateral diffusion rates of DLPC lipids were calculated from both polymer- and MSP-nanodiscs from the end 0.5 μs MD simulation.

MD trajectories were interpreted using visual molecular dynamics^[49]^ and images were built using PyMOL (academic license, https://pymol.org/2/) and Discovery studio visualizer 3.5 (Accelrys).^[50]^ All CG MD simulations were carried out using Martini v 2.2P^[24]^ force field in GROMACS MD engine running parallel in SGI UV 3000. The list of MD simulation parameters are given in Table-S1.

### 4.6 Simulated annealing CG-MD

The thermal stability of PMAQA- or SMAEA-DLPC nanodiscs were tested by performing simulated annealing CG-MD simulation by linearly increasing temperature over time as described elsewhere^[51]^. A total of 14 annealing points were considered with coupling temperatures ranging from 298 K up to 353 K on a time scale of 10 μs. The initial group was coupled to 298 K and the MD system was linearly heated up (increased in every 2 μs up to 6 μs). A constant temperature of 353 K was applied to both polymer-nanodisc systems from 6 to 10 μs to monitor nanodisc destabilization. The MD snapshots were retrieved in every 2 μs and superimposed using DSV. A reference atomic distance of 7.5 Å was defined in DSV and the nanodisc image was exported to ImageJ (NIH). The Feret’s diameter, area and perimeter of the discoidal shaped nanodisc simulated at different temperature points were next analyzed using *ImageJ* and plotted in *Origin.*

## Acknowledgments

This study was supported by funds from NIH (AG048934 to A.R.). We thank Dr. Charles L. Brooks III for fruitful discussion on the use of molecular dynamics simulations to study nanodiscs. This work was (in part) performed under the International Collaborative Research Program of Institute for Protein Research, Osaka University, ICR-18-02. We thank Professor Toshimichi Fujiwara in the Institute for Protein Research, Osaka University, for providing parallel computing facility on *SGI* UV 3000.

