## Supplemental Data for "Self-assembly of polymer-encased lipid nanodiscs and membrane protein reconstitution"

**Table S1.** Parameters of CG MD simulation systems

| Polymer (number) | Lipid | Protein (Number) | Water | Ions | Time (μs) |
|---|---|---|---|---|---|
| SMAEA (50) | DLPC (200) | - | 40174 | 450 | 10 |
| SMAEA (15) | DLPC (60) | - | 11295 | 135 | 10 |
| PMAQA* (15) | DLPC (60) | - | 11641 | 90 | 10 |
| PMAQA (8) | DLPC (64) | - | 11455 | 120 | 10 |
| PMAQA (15) | DLPC (60) | - | 10863 | 225 | 10 |
| SMAEA (15) | DLPS (60) | - | 11203 | 195 | 10 |
| PMAQA (15) | DLPS (60) | - | 11683 | 30 | 10 |
| SMAEA (15) | DLPC (60) | srII (1) | 11295 | 135 | 10 |
| SMAEA (15) | DLPC (60) | APP (1) | 11291 | 139 | 10 |
| SMAEA (15) | DLPC (60) | Integrin-β3 (1) | 11294 | 136 | 10 |

* denotes ~2.0 kDa PMAQA

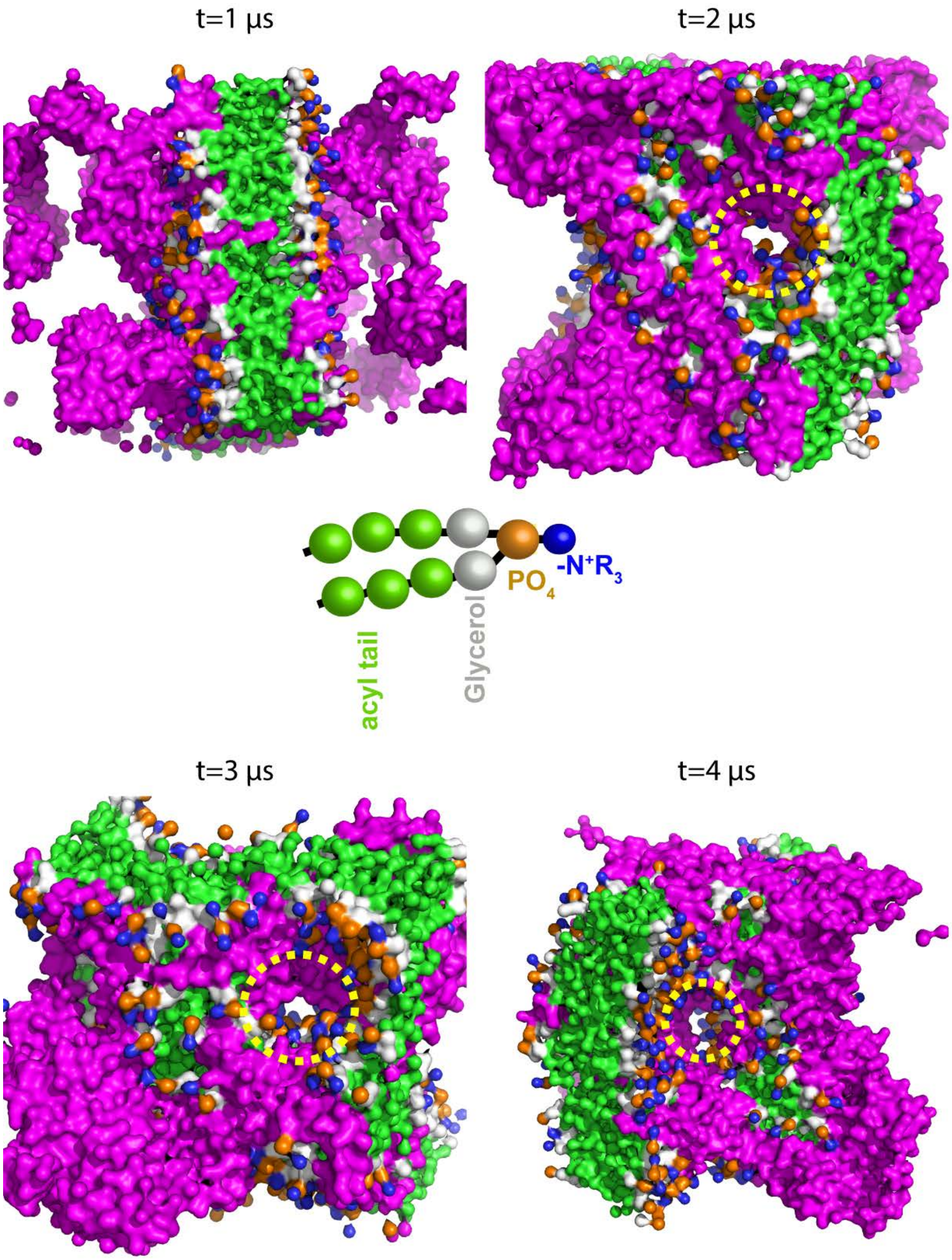


**Figure S1.** MD snapshots showing interaction of SMAEA with DLPC lipid bilayer at different time scale. The complex structure is presented as surface in PyMOL. The coarse-grained model of SMAEA molecules are shown in pink and lipid groups are indicated in different colors as shown in the center. The SMAEA insertion and formation of pore in DLPC lipid-bilayer is shown inside a yellow dashed circle. The water and ions are not shown for clarity.

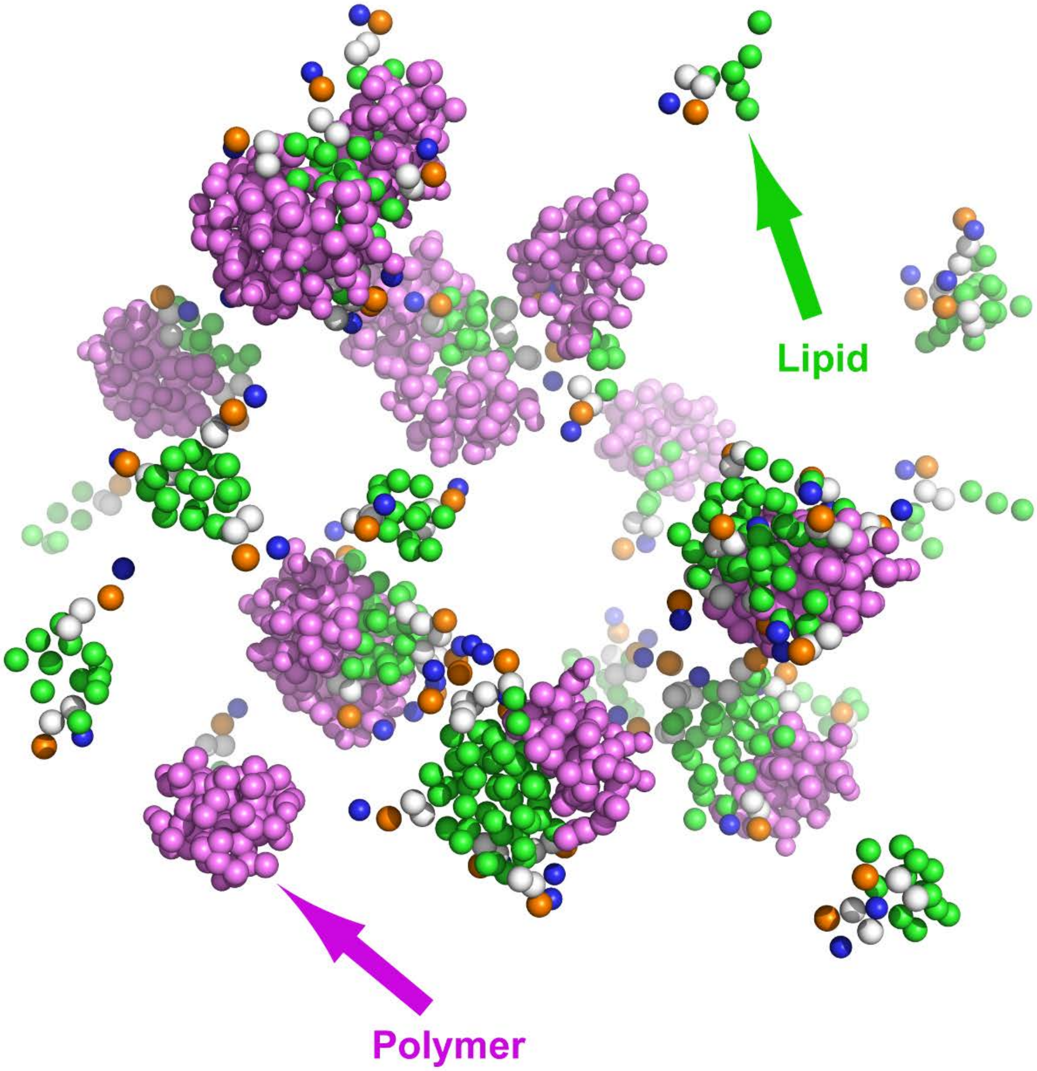


**Figure S2.** Random distribution of copolymers and DLPC lipids in solvent for spontaneous nanodisc generation. The coarse-grained model of SMAEA molecules are shown in pink and lipid groups in different colors as indicated in Figure. S1 (center). The water and ions are not shown for clarity.

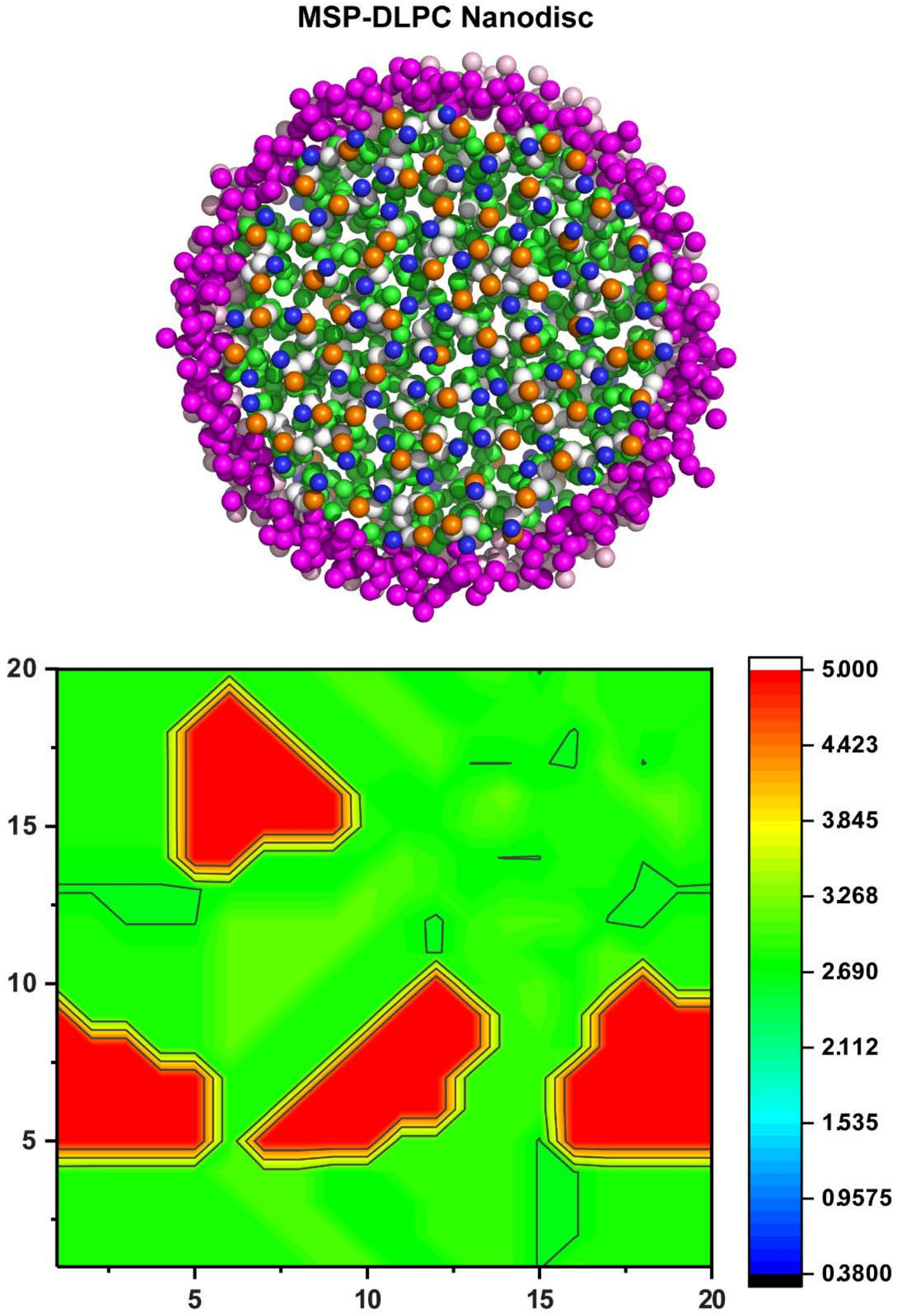


**Figure S3.** Coarse-grained model of membrane scaffold protein (MSP) encased DLPC nanodisc generated using CHARMM_GUI[16] (A). The MSP protein chains are shown in light and dsark pink and the lipid groups in different colors as indicated in Figure. S1 (center). (B) Lipid-bilayer thickness of MSP-DLPC nanodisc calculated using GridMAT-MD[15] from a 20x20 matrix. The scale for the bilayer thickness in nanometer is shown on the right. The water and ions are not shown for clarity.

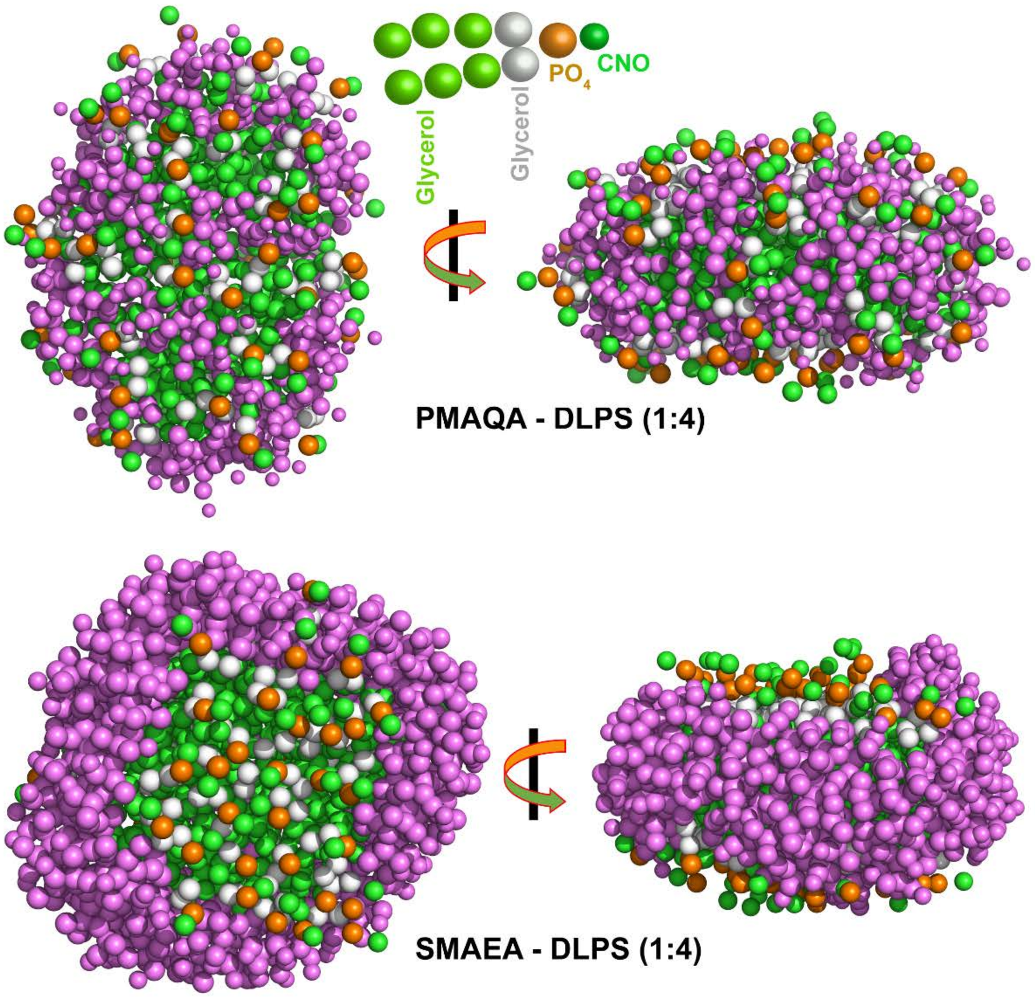


**Figure S4.** MD snapshots showing formation of DLPS nanodiscs by PMAQA and SMAEA at the indicated polymer to DLPS molar ratio. Coarse-grained models of DLPS nanodiscs retrieved at 10 μs are shown in top and side-view for both PMAQA and SMAEA. The polymers are shown in pink and lipid groups are shown on the top. The water and ions are not shown for clarity.

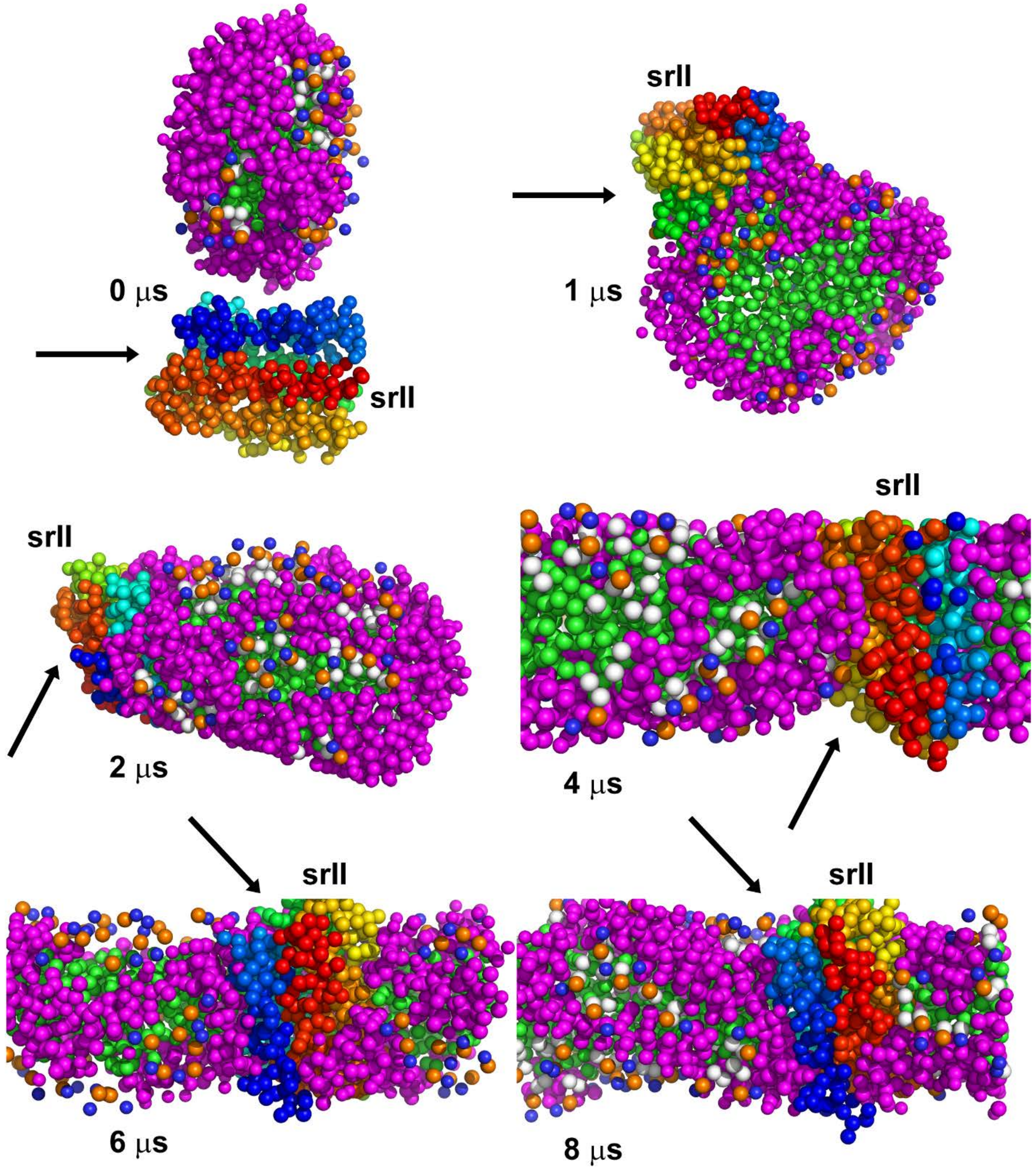


**Figure S5.** MD snapshots showing interaction or reconstitution of seven transmembrane domain bacterial sensory rhodopsin (srII) in SMAEA-DLPC nanodisc at different time scale. The protein molecule is color as spectrum in PyMOL with N-terminal in blue and C-terminal in red. SMAEA molecules are shown in pink and lipid groups in different colors as indicated in Figure.S1 (center). The arrows indicate the binding of srII to nanodiscs and its translational motion along the bilayer normal as a function of time. The water and ions are not shown for clarity.

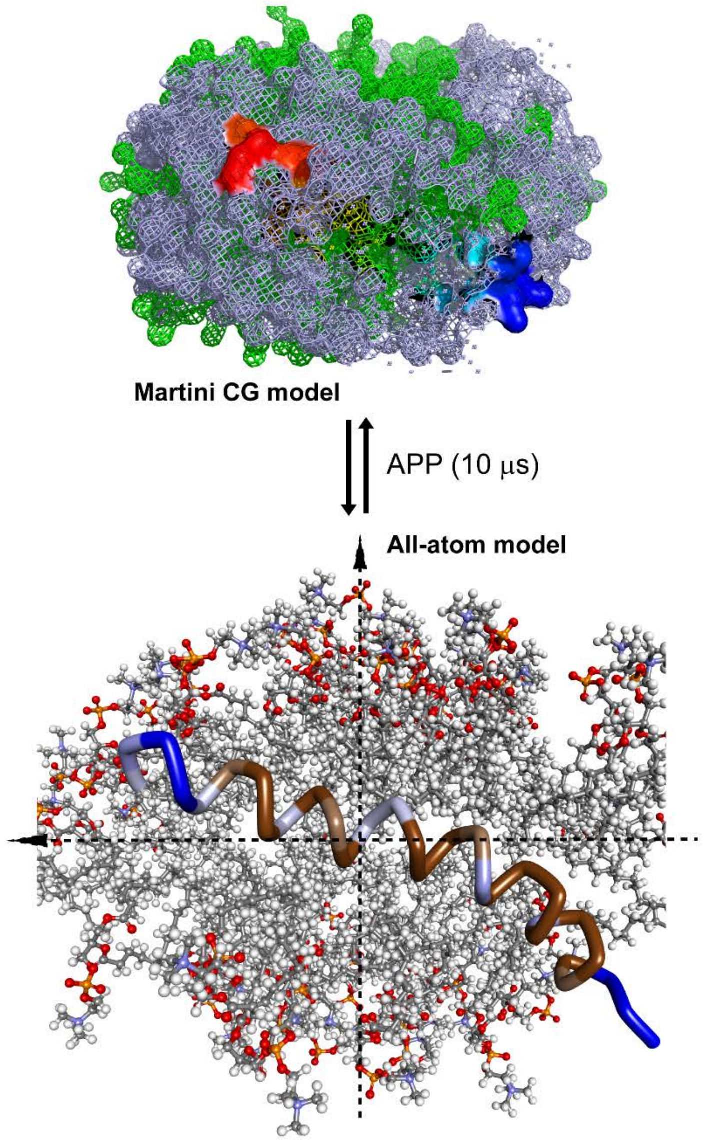


**Figure S6.** A representative coarse-grained (CG) to all-atom converted model structure of single transmembrane domain amyloid-precursor protein (APP) bound to SMAEA-DLPC nanodisc and retrieved at 10 μs from MD simulation. The APP molecule is color as spectrum in PyMOL with N-terminal in blue and C-terminal in red in the CG model structure (top). SMAEA and lipid molecules are shown in mesh as grey and green, respectively. The all-atom model of APP is shown as tube and DLPC in CPK in Discovery studio visualizer. The dashed line indicate a tilt angle of ≈ 20° in APP with respect to the plane of bilayer. The water and ions are not shown for clarity.

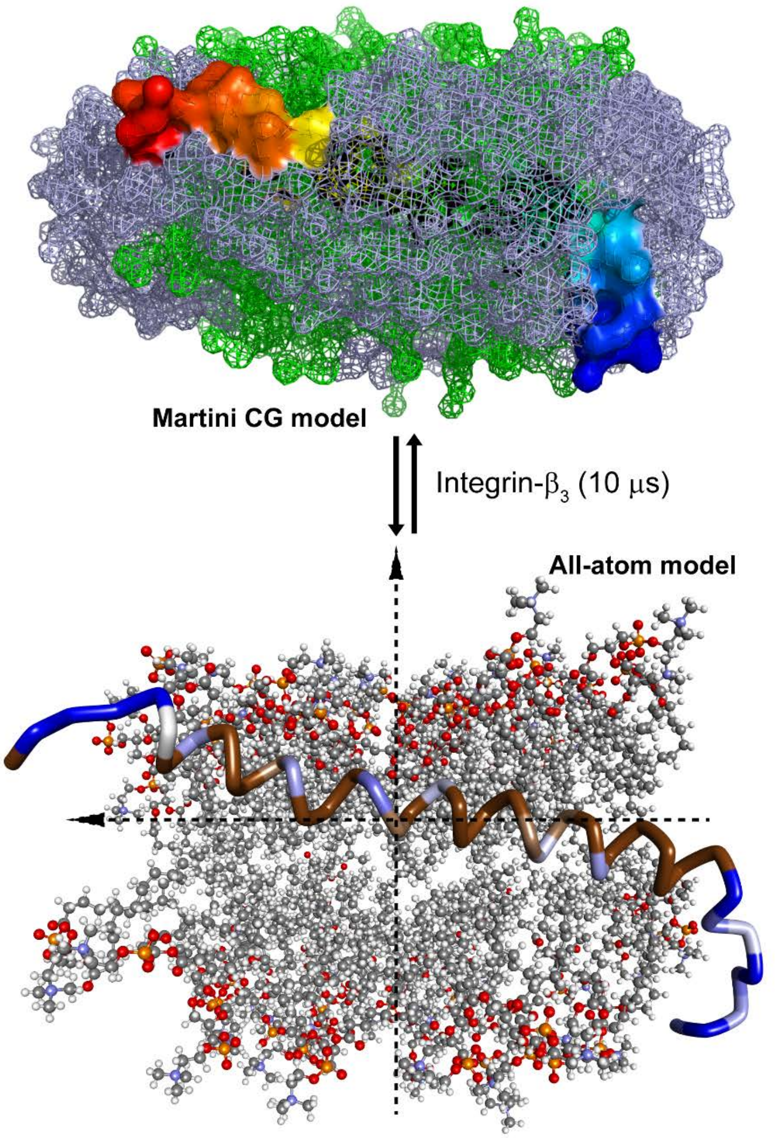


**Figure S7.** A representative coarse-grained (CG) to all-atom converted model structure of integrin-β3 single transmembrane domain protein constituted in SMAEA-DLPC nanodisc and retrieved at 10 μs from MD simulation. The integrin-β3 molecule is color as spectrum in PyMOL with N-terminal in blue and C-terminal in red in the CG model structure (top). SMAEA and lipid molecules are shown in mesh as grey and green, respectively. The all-atom model of integrin-β3 is shown as tube and DLPC in ball-stick in Discovery studio visualizer. The dashed line indicate a tilt angle of ≈30° in integrin-β3 with respect to the plane of bilayer. The water and ions are not shown for clarity.
